## Supplemental information for "Single-cell RNA sequencing identifies phenotypically, functionally, and anatomically distinct stromal niche populations in human bone marrow"

Figure S1

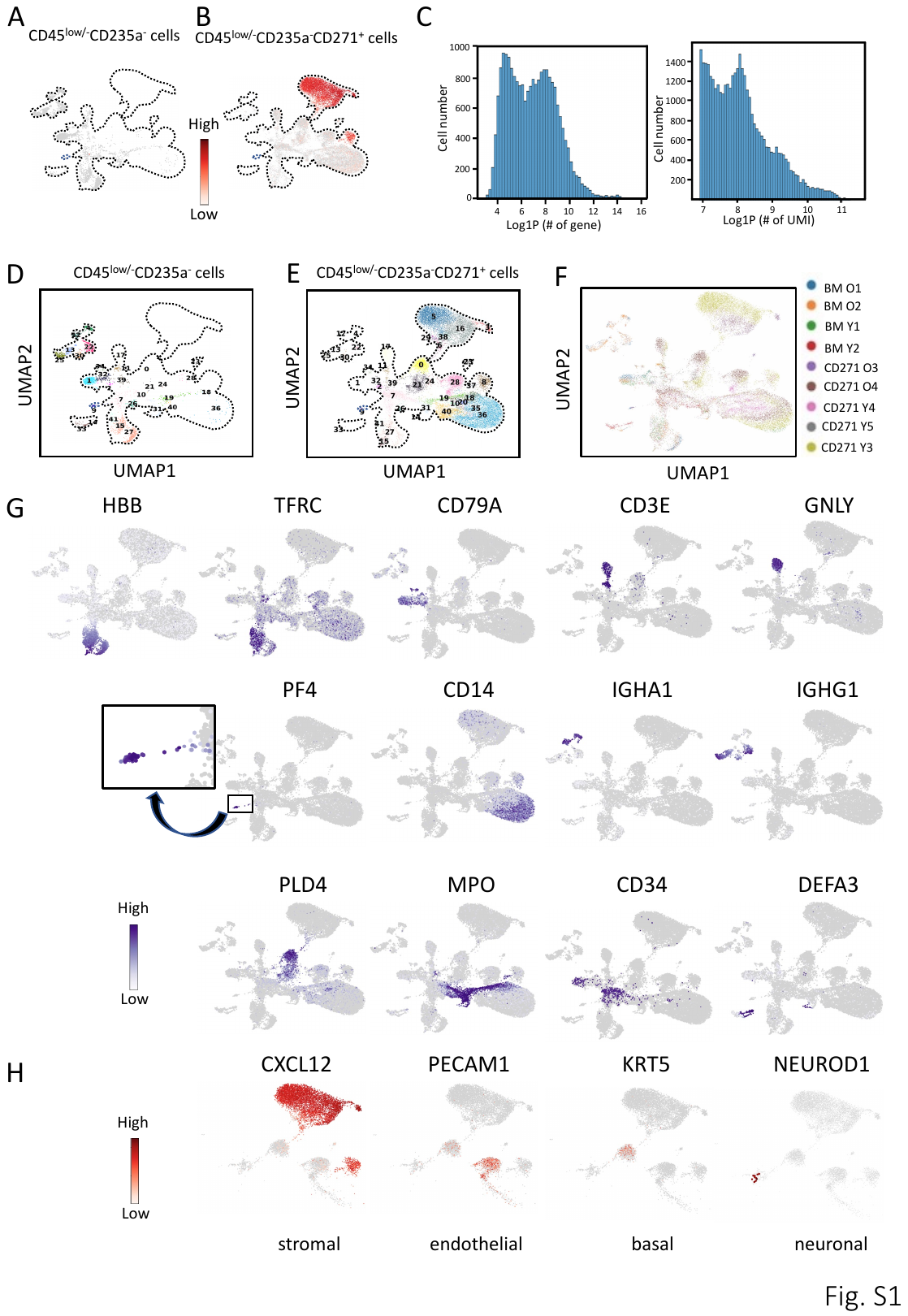

Figure S1. UMAP display of normalized CXCL12 expression in sorted CD45^low/-^CD235a^-^ cells (5704 cells) (A) and CD45^low/-^CD235a^-^CD271^+^ cells (19363 cells) (B). Dashed lines outline shows the UMAP overview of the entire dataset. (C) scRNAseq quality measures expressed as gene numbers (left panel) and unique molecular identifier (UMI, right panel) versus cell number (y-axis). Cells with less than 1000 UMIs were removed. (D) UMAP display of single-cell transcriptomic data of human bone marrow CD45^low/-^CD235a^-^ cells (5704 cells) from four healthy donors. Cluster number and color legend are consistent with Figure 1B. (E) UMAP display of single-cell transcriptomic data of human bone marrow CD45^low/-^CD235a^-^CD271^+^ cells (19363 cells) from five healthy donors. Cluster number and color legend are consistent with Figure 1B. (F) UMAP display (as in Fig. 1B) of human bone marrow CD45^low/-^CD235a^-^ cells and CD45^low/-^CD235a^-^CD271^+^ cells from nine healthy donors with samples color coded (25067 cells). BM indicates CD45^low/-^CD235a^-^ cell donors (n=4), CD271 indicates CD45^low/-^CD235a^-^CD271^+^ cell donors (n=5). O, old; Y, young (see also Materials and Methods). (G) UMAP displays (as in Fig. 1B) highlighting the expression of selected hematopoietic signature genes for major cell types. PF4-expressing cells were blown-up for better visualization. (H) UMAP display (as in Fig. 1C) of normalized expression of selected non-hematopoietic signature genes for major cell types. Expression was normalized to 1000 UMIs. Scale bars are adjusted for optimal visualization in (G) and (H).

Figure S2

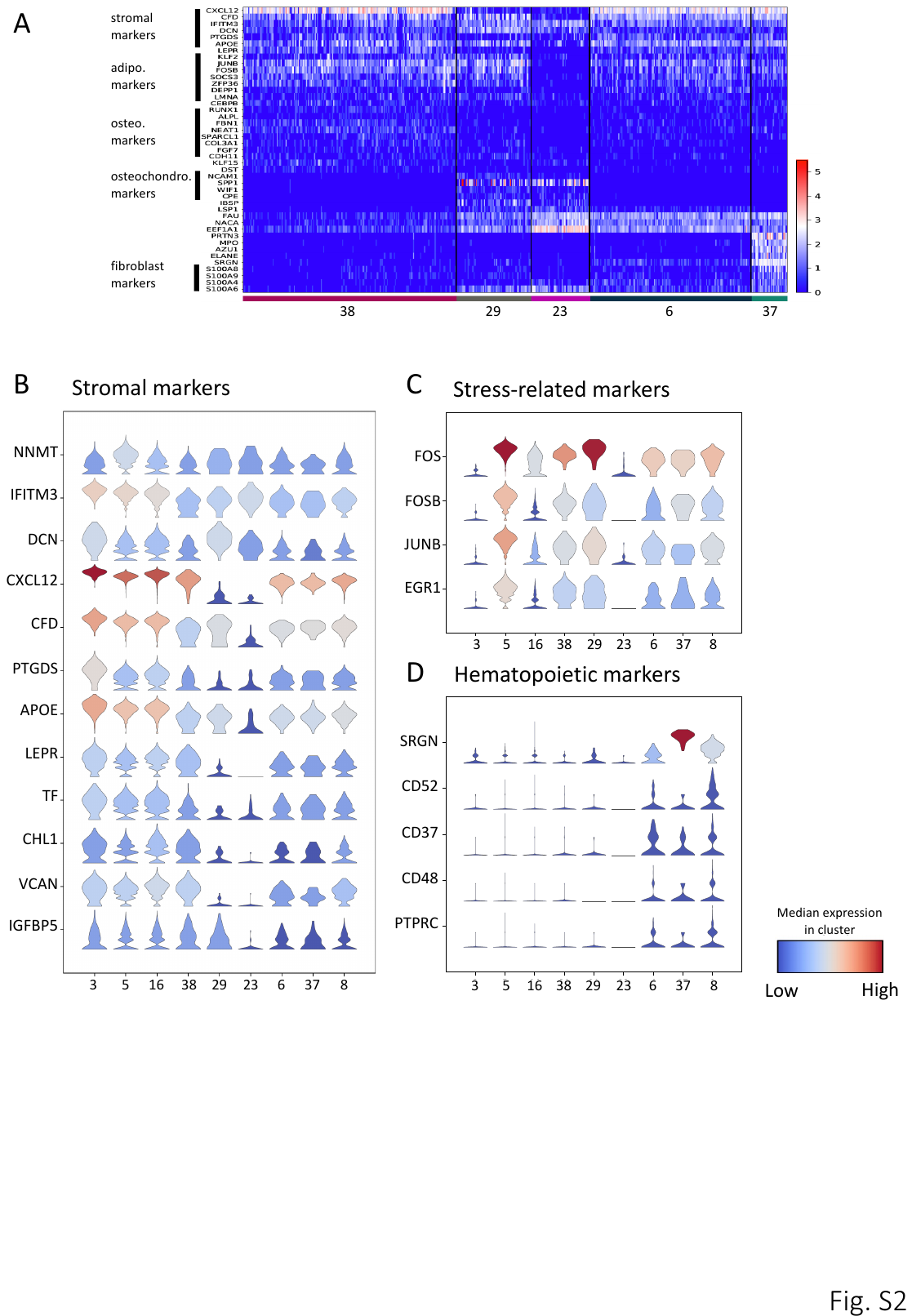

Figure S2. (A) Single cell heatmap of representative differentially expressed genes in clusters 38, 29, 23, 6 and 37. Stacked violin plots of stromal markers (B), stress-related transcription factors (C) and hematopoietic markers expressed by stromal clusters (D). Cluster annotation as in Fig 2A.

Figure S3

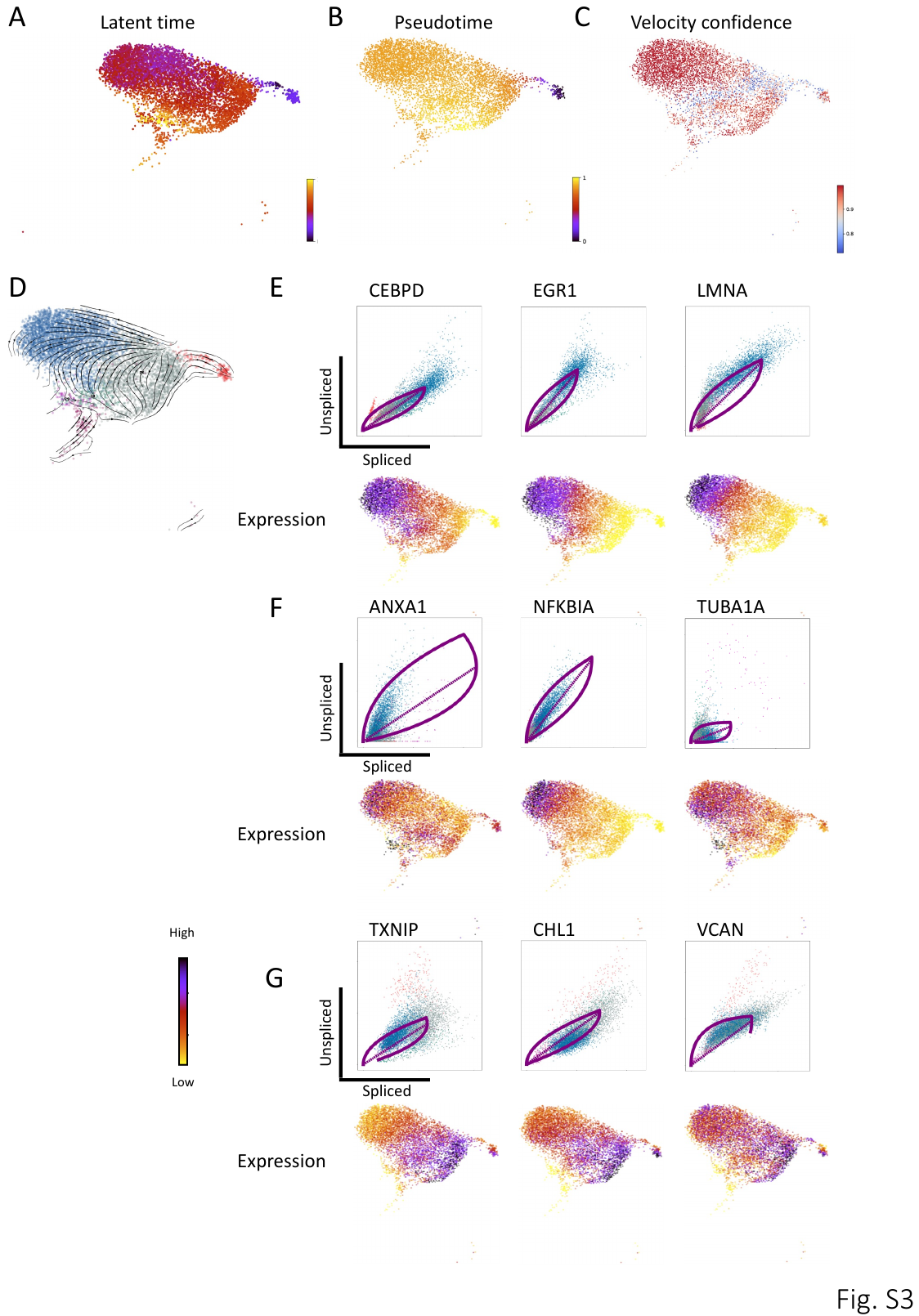

Figure S3. (A-B) UMAP display of latent time and pseudotime analyses for the selected stromal clusters that formed a continuum of different cellular states. Inferred latent time and pseudotime are represented by a color scale from 0 (the earliest latent time/pseudotime) to 1 (the latest latent time/pseudotime). (C) UMAP display of velocity confidence for the selected stromal cells. Scale bar indicates the average correlation of the velocity vector of a certain cell and those of its neighbors. (D) Different cellular states of the selected stromal cells were analyzed with scVelo and single cell velocities visualized as streamlines in a UMAP. Black arrows indicate direction and thickness indicates speed along the stromal cell development trajectory. Colors correspond to cluster colors in Figure 3C. (E-G) Expression dynamics of putative driver genes identified by likelihood model analysis. Upper panel: Phase portraits of selected putative driver genes (indicated on top) characterize their splicing kinetics. The dashed purple line corresponds to the estimated ‘steady-state’ ratio, i.e. the ratio of unspliced to spliced mRNA abundance. Positive (cells over the dashed purple line) and negative velocities (cells under the dashed purple line) correlate with up- and down-regulation of genes, respectively. The solid purple lines indicate the learned kinetics for each gene calculated with a likelihood-based model by scVelo. The colors in the phase portraits correspond with the colors in UMAP (as in Fig. S3D). Lower panel: UMAP display (as in Fig. 3C) of the expression dynamics of putative driver genes indicated on top of the phase portraits in the upper panel. Colors correlate with expression levels as shown in the color bar.

Figure S4

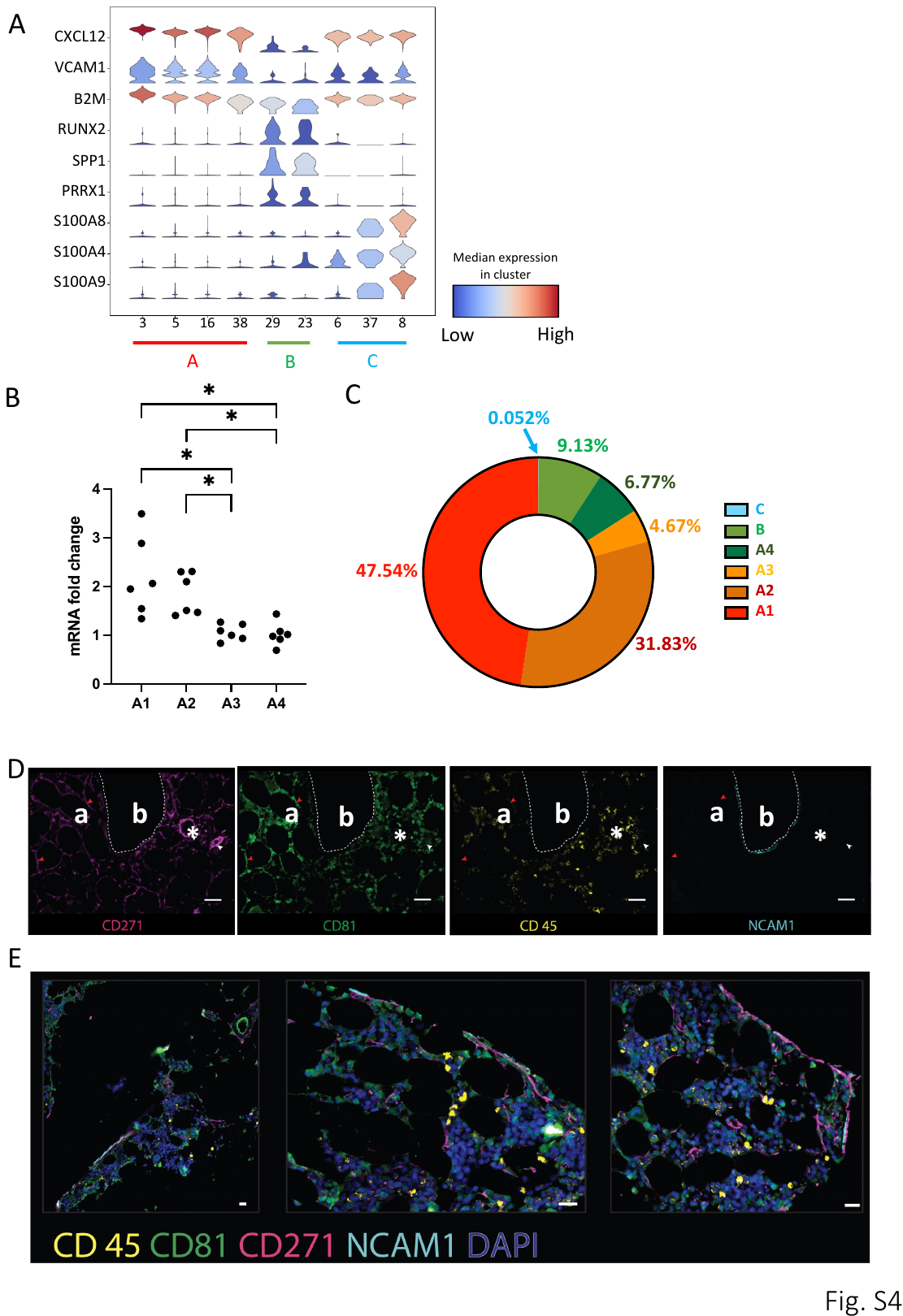

Figure S4. (A) Stacked violin plots of stromal, osteochondrogenic, and fibroblastic gene expression in different stromal clusters. Corresponding stromal cell groups are indicated on the x-axis legend. The color bar indicates gene expression in each cluster. (B) mRNA fold change of CD81 in FACS soreted A1-A4 subsets (n=3). Results are shown as mRNA fold change after standardization with GAPDH levels. For comparison, expression level of CD81 in A4 is set as one fold. *: p<0.05. (C) The relative contribution of each sorted stromal cell populations in (B) to total CFU-F. Data are given as a normalized mean percentage as indicated in the figure (n=3). The sum of CFU-F of all populations is defined as 100%. (D) Formalin-fixed, paraffin-embedded (FFPE) human BM slides were sequentially stained for CD271 (pink), CD81 (green), NCAM1 (cyan) and CD45 (yellow) and scanned with the OlympusVS120 slide scanner. Single staining for each marker is shown as indicated under each picture. Scale bars represent 50 µm. Red arrows: CD271^+^CD81^++^ cells; white arrows: arteriolar walls; white lines: bone lining regions. Bone (b), adipocytes (a), and capillaries (*) are indicated. (E) FFPE human BM slides were sequentially stained for DAPI (blue), CD45 (yellow), CD81 (green), CD271 (pink), and NCAM1 (cyan) and scanned with the OlympusVS120 slide scanner. Scale bars represent 20 µm.

Figure S5

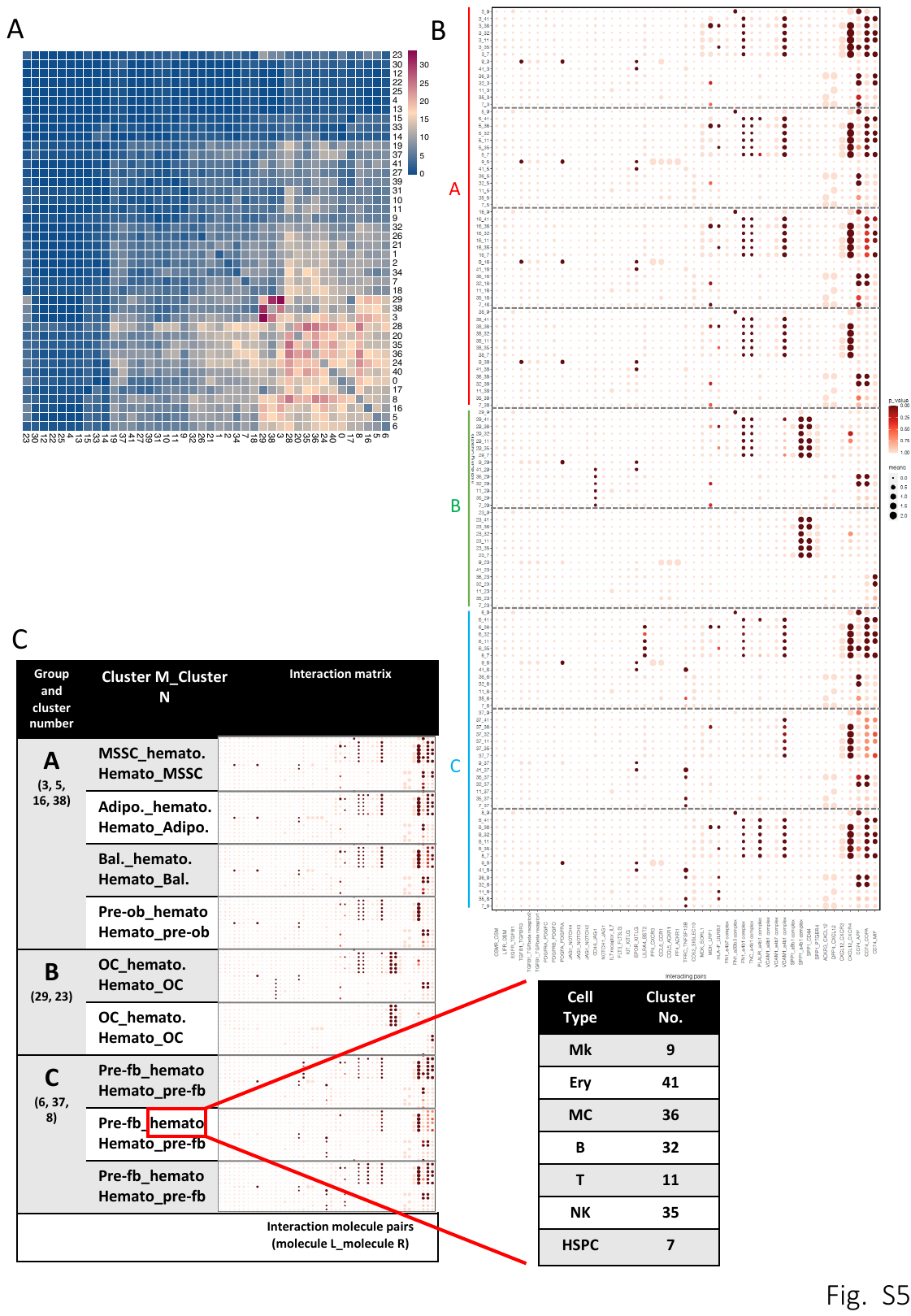

Figure S5. (A) Heatmap showing the total number of interactions between different clusters in bone marrow CD45^low/-^CD235a^-^ cells. Scale bar colors represent numbers of interactions. (B) Dot plot of selected ligand-receptor pairs between bone marrow stromal cells and different hematopoietic cell types. The y-axis label indicates the pair of interacting cell clusters (‘cluster X_cluster Y’ indicates that cluster X interacts with cluster Y by ligand-receptor expression). The x-axis indicates the interacting receptor/ligand (R/L) or ligand/receptor pairs, respectively (‘molecule L_molecule R’ means molecule L interacts with molecule R). The means of the average expression level of the interacting molecules are indicated by circle size. The scale is shown on the right side. P values are indicated by circle color and correspond to the adjacent upper scale bar. Dashed lines separate the interactions between hematopoietic cells and different stromal clusters. (C) Schematic overview of the layout of the interaction matrix in (B). Adipo., adipo-primed progenitors; Bal., balanced progenitors; pre-ob, pre-osteoblasts; hemato, hematopoietic clusters. Cluster numbers of the hematopoietic cell types selected for this analysis are listed in the table on the right side.

Figure S6

Figure S6. (A-E, G-H, J) UMAP (as in Fig. 1B) illustration of the normalized expression of selected ligand and receptor gene pairs. (F) Flow cytometric analysis of SPP1 expression of primary CD45^low/-^CD235a^-^CD71^-^CD271^+^CD56^+^ cells (red histogram), CD45^low/-^CD235a^-^CD71^-^CD271^+^CD56^-^CD81^++^ cells (blue histogram) and corresponding isotype control (orange histogram). (I) CFU-F assay of sorted CD45^low/-^CD235a^-^CD71^-^CD271^+^ cells (100 cells/well) in the presence (upper panel) or absence (lower panel) of sorted bone marrow CD45^+^ cells (3x10^5^ cells/well), respectively.

Figure S7

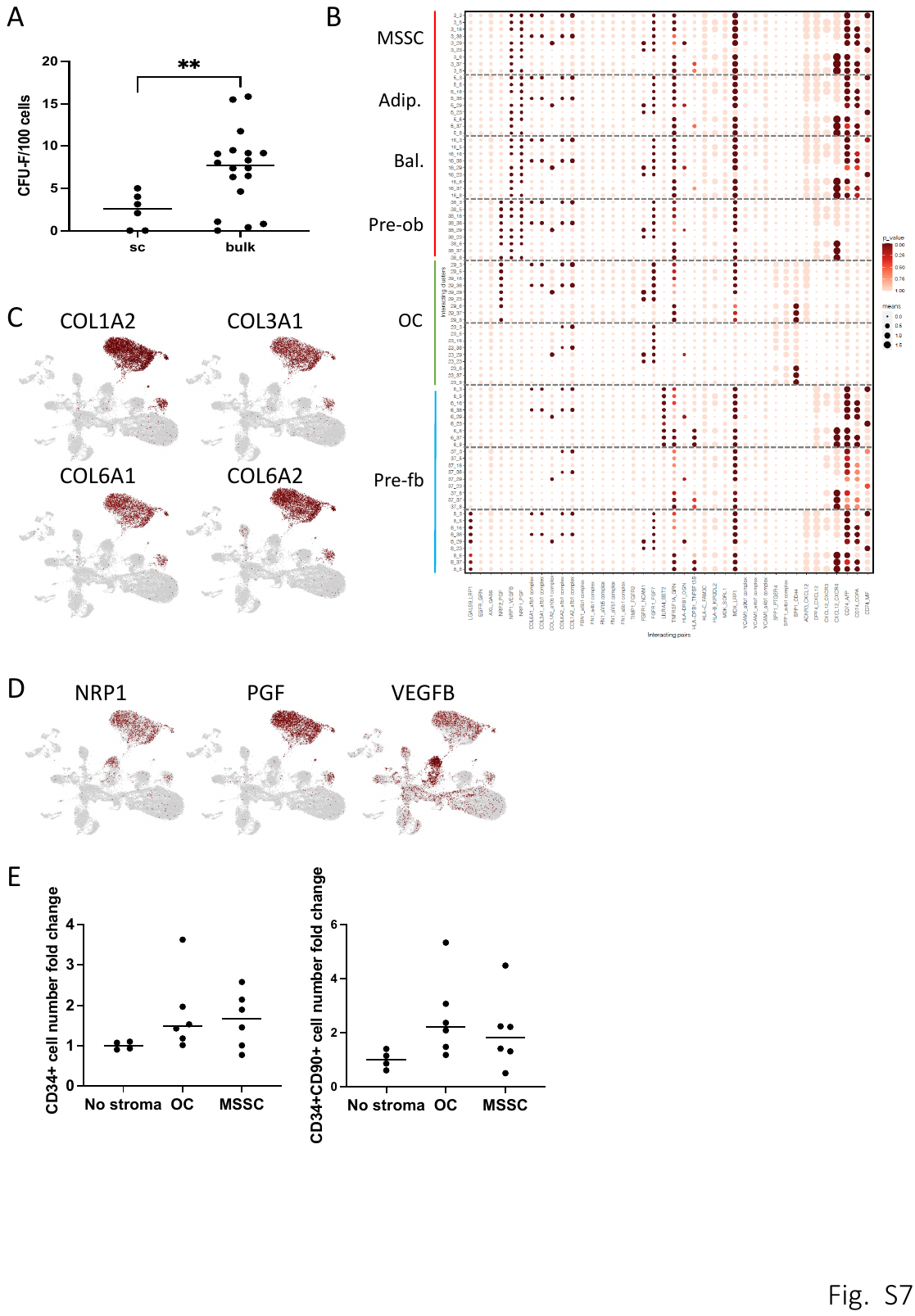

Figure S7. (A) CFU-F frequency of single-cell and bulk sorted MSSCs, i.e. sorted A1 cells (CD45^low/-^CD235a^-^CD71^-^CD271^+^NCAM1^-^CD52^-^CD81^++^). Data are shown as individual data points and medians (horizontal lines), n=3. **: p<0.01. (B) A dot plot overview of top-ranked ligand-receptor pairs between different bone marrow stromal clusters. P values are indicated by circle color. The means of the average expression levels are indicated by circle size. Scale bars are provided on the right side of the plot. Stromal cell cluster pairs are indicated by “cluster number_cluster number” (y-axis labels). Dashed lines separate the different stromal clusters. Adipo., adipo-primed progenitors; Bal., balanced progenitors; Pre-ob, pre-osteoblasts; Pre-fb, pre-fibroblasts. Interacting molecule pairs are indicated in the x-axis labels. (C) UMAP (as in Fig. 1B) illustration of the normalized expression of genes involved in alpha 1 beta 1 integrin complex (a1b1 complex). (D) UMAP (as in Fig. 1B) illustration of the normalized expression of selected ligand and receptor genes. (E) Fold change of total number of CD34^+^ cells (left) and CD34^+^CD90^+^ cells (right) produced after seven days in culture. Results are shown as fold change relative to the cell number of standard CD34^+^ culture without stroma support (No stroma). Data are presented as individual data (dots) and median (horizontal lines) from independent experiments (n=2).

**List of supplemental tables**

**Table S1. Donor information (related to Fig. 1A).**

Sample ID, age, gender and selected cell population for single cell RNA sequencing were listed.

**Table S2. Cluster annotation (related to Fig. 1B).**

Cluster ID, annotation and marker gene(s) used for annotation were listed.

**Table S3. Cell number and percentage of each cluster in each sample.**

Table S3. Sheet 1. Cell number and percentage of each cluster in each sample (related to Fig. 1B).

Table S3. Sheet 2. Cell numbers and percentages of stromal clusters in each sample (related to Fig. 2).

Table S3. Sheet 3. Cell numbers and percentages of non-hematopoietic clusters in each sample (related to Fig. 1C).

**Table S4. DE (differentially expressed) genes in each cluster (related to Fig. 1B).**

DE genes for each cluster were listed in the individual sheet.

**Table S5. List of the 300 top-ranked likelihood genes shown in Figure 3C.**

**Table S6. List of ligand-receptor interactions in scRNAseq dataset as identified by CellPhoneDB analysis (related to Fig. 5, S5 and Fig. 7A).**

Means (measurement for interaction strength), p values and significant p value for each interaction are listed in individual sheets. In the first row of each sheet, X|Y indicates the interaction between cluster X and cluster Y. The empty spaces in the sheet ‘significant_means’ indicate that non-significant means were removed.
