## Supplementary material for "Single-cell RNA sequencing identifies phenotypically, functionally, and anatomically distinct stromal niche populations in human bone marrow": Table S1 and S2

Table S1. Donor information

| Sample ID | Age | Gender | Cell population |
| --- | --- | --- | --- |
| BM_Y1 | 19 | F | CD45^low/-^CD235a^-^ |
| BM_Y2 | 22 | M | CD45^low/-^CD235a^-^ |
| BM_O1 | 52 | M | CD45^low/-^CD235a^-^ |
| BM_O2 | 53 | F | CD45^low/-^CD235a^-^ |
| CD271_Y3 | 21 | F | CD45^low/-^CD235a^-^CD271^+^ |
| CD271_Y4 | 25 | M | CD45^low/-^CD235a^-^CD271^+^ |
| CD271_Y5 | 32 | M | CD45^low/-^CD235a^-^CD271^+^ |
| CD271_O3 | 58 | F | CD45^low/-^CD235a^-^CD271^+^ |
| CD271_O4 | 61 | M | CD45^low/-^CD235a^-^CD271^+^ |

Table S2. Cluster annotation

| Cluster ID | Annotation | Marker gene(s) |
| --- | --- | --- |
| 0 | KRT5-enriched basal cells | KRT5 |
| 1 | B-cell progenitors | DNTT, VPREB1, VPREB3, CD79A, CD79B, IGLL1 |
| 2 | B-cell progenitors | DNTT, VPREB1, VPREB3, CD79A, CD79B, IGLL1 |
| 3 | Stromal cells | CXCL12, VCAN, LEPR |
| 4 | Plasma cells | IGHA1, IGHA2, IGKC |
| 5 | Stromal cells | CXCL12, VCAN, LEPR |
| 6 | Stromal cells | CXCL12, VCAN, LEPR |
| 7 | HSPC/CD34-enriched | CD34, PROM1 (CD133), CRHBP, AVP, MLLT3, FAM30A, GATA1 |
| 8 | Stromal cells | CXCL12, VCAN, LEPR |
| 9 | Megakaryocytes | PF4, GP9, PPBP, PPBPP2 |
| 10 | Dendritic cells | FLT3, PLAC8, PLD4, GZMB, IRF7, IRF8, NAPSB |
| 11 | T cells | IL7R, CD3D, CD3E, CD3G |
| 12 | Plasma cells | IGKC, IGHA1, IGHA2, IGLC3 |
| 13 | Plasma cells | IGKC, IGHG1, IGHG2, IGHG3, IGHG4, IGHGP |
| 14 | Granulocytic cells | DEFA3, DEFA4 |
| 15 | Erythroid cells | HBB, HBA1, HBA2, AHSP, TFRC |
| 16 | Stromal cells | CXCL12, VCAN, LEPR |
| 17 | NK cells | NCAM1, GZMH, GNLY, GZMA, IL32 |
| 18 | Granulocytic cells | MPO, SRGN |
| 19 | Granulocytic cells | MPO, AZU1, PRTN3, SRGN, ELANE |
| 20 | Dendritic cells | PLAC8, GZMB, IRF8, PLD4 |
| 21 | Dendritic cells | PLAC8, STMN1, IRF8 |
| 22 | Plasma cells | IGLC2, IGHG1, IGHG3, IGLC3, IGHGP, IGKC |
| 23 | Stromal cells | CXCL12, VCAN, LEPR |
| 24 | Dendritic cells | SCT, CST3, PLAC8 |
| 25 | Plasma cells | IGHG1, IGHGP, IGHG3, IGKC, IGHG4 |
| 26 | Granulocytic cells | CLC, SRGN, MS4A3, ANXA1 |
| 27 | Erythroid cells | HBB, HBA1, HBA2, AHSP, TFRC |
| 28 | Endothelial cells | PECAM1, ICAM2 |
| 29 | Stromal cells | CXCL12, VCAN, LEPR |
| 30 | Plasma cells | IGKC, IGHG3, IGHG1 |
| 31 | Granulocytic cells | AZU1, ELANE, MPO, SRGN |
| 32 | B-cell progenitors | DNTT, VPREB1, VPREB3, CD79A, CD79B, IGLL1 |
| 33 | Granulocytic cells | CEACAM8, PGLYRP1, TCN1 |
| 34 | B-cell progenitors | CD79A, CD79B |
| 35 | NK cells | NCAM1, GZMH, GNLY, GZMA, GZMB, IL32 |
| 36 | Monocytes | CSF1R, CD14, CD33, ITGAM (CD11B), CD86 |
| 37 | Stromal cells | CXCL12, VCAN, LEPR |
| 38 | Stromal cells | CXCL12, VCAN, LEPR |
| 39 | Neuron-enriched | NEUROD1, CHGB, ELAVL4, STMN2, INSM1 |
| 40 | Dendritic cells | NAPSB, FLT3, IRF8, PLD4, IRF7 |
| 41 | Erythroid cells | HBB, HBA1, HBA2, AHSP, TFRC |

Abbreviations: HSPC, hematopoietic stem/progenitor cells; NK, natural killer
